# Environmentally mediated reproductive success predicts breeding dispersal decisions in an early successional amphibian

**DOI:** 10.1101/349621

**Authors:** Laurent Boualit, Julian Pichenot, Aurélien Besnard, Rémi Helder, Pierre Joly, Hugo Cayuela

## Abstract

Dispersal is a central mechanism in ecology and evolution. Dispersal evolution is driven by a trade-off between costs and benefits, which is influenced by inter-individual variability and local environmental conditions (context-dependent dispersal). Many studies have investigated how dispersal decisions may be influenced by environmental factors, including density, predation, and interspecific competition. Yet few have attempted to examine how habitat disturbance may affect the dispersal process in spatially structured populations. In early successional species, one might expect individuals to adjust their dispersal decisions based on two main factors that potentially have an influence on reproductive success: patch size and the level of patch disturbance. In this study, we examined how these two factors affect breeding success and dispersal decisions in an early successional amphibian, the yellow-bellied toad (*Bombina variegata*). To this end, we used capture–recapture data collected on a spatially structured population occupying 28 breeding patches. We took advantage of recent developments in multievent capture–recapture models to detect signs of context-dependent dispersal. The results revealed that the probability of successful reproduction and the number of newly metamorphosed individuals increased with both the size and the proportion of disturbance of a patch. In addition, our results showed that the factors affecting breeding success also influenced breeding dispersal probability. Large patch size negatively influenced emigration probability; in contrast, it positively influenced immigration probability. Equally, higher disturbance (in terms of the proportion of the patch’s surface area disturbed each year) had a strong negative influence on emigration probability and slightly positively affected immigration probability. These findings strongly suggest that individuals make context-dependent dispersal decisions, adjusted to maximize future fitness prospects in a patch, allowing them to better cope with rapid changes in environmental conditions resulting from the ecological succession process. This opens new areas of potential research into the role of dispersal in organism specialization along an ecological succession gradient.

## Introduction

Dispersal, the movement of an individual from its site of birth to its reproduction site (i.e. natal dispersal), or between successive reproduction sites (i.e. breeding dispersal), is a central mechanism in ecology and evolution (Ronce 2007, Clobert et al. 2009, Matthysen 2012). Dispersal has a broad influence on the dynamics of spatially structured populations, as it affects local population density, the risk of local extinction, and the possibility of patch (re)colonization (Hanski & Gaggiotti 2004, Gilpin 2012). Moreover, it has a strong influence on evolutionary processes since it affects genetic variation and adaptation through gene flow (Ronce 2007, Legrand et al. 2017). Dispersal is usually considered a three-stage process that includes emigration (or departure), transience (or transfer within the landscape matrix) and immigration (or arrival) (Ims & Yoccoz 1997, Ronce 2007). A major advance in dispersal studies was the recognition that dispersal evolution is driven by a balance between costs and benefits at each stage of the process (Bonte et al. 2012). This trade-off is influenced by factors related to the individual (phenotype-dependent dispersal) and to local environmental conditions (context-dependent dispersal) (Clobert et al. 2009, Matthysen 2012). Contrary to the assumption made in many demographic models, dispersal is thus not a random process (Edelaar et al. 2008). Individuals adjust their dispersal decisions according to environmental and social cues that provide information about their future fitness prospects in a given patch (i.e. informed dispersal, Clobert et al. 2009), resulting in asymmetric dispersal rates in spatially structured populations.

Many studies have investigated how dispersal decisions may be influenced by environmental factors such as density, predation, interspecific competition and landscape characteristics (reviewed in Bowler & Benton 2005, Matthysen 2012, Cote et al. 2017). Yet few have attempted to examine how the level of habitat disturbance may affect the dispersal process in spatially structured populations (Bates et al. 2006, Altermatt & Ebert 2008, 2010, Duckworth 2012). A disturbance is a temporary change in physical environmental conditions (e.g. due to fire, flood or drought) and can be caused by natural or anthropogenic factors. Disturbances play a central role in community and ecosystem dynamics by initiating ecological successions, i.e. the sequential replacement of species following the loss of biomass due to a disturbance event (Turner et al. 1998, Prach & Walker 2011). In addition, disturbances can create new habitat patches by reshaping the physical environment. In early successional species (i.e. those occurring at the early stages of succession), the disturbance regime has a strong influence on population dynamics, as it affects the distribution and the amount of suitable habitat across a landscape (Moloney & Levin 1996, Amarasekare & Possingham 2001, Meurant 2012). In this context, dispersal is expected to be a central mechanism in population dynamics, as it allows individuals to escape rapid detrimental changes in environmental conditions (e.g. declining quality of a patch through the succession process) and to colonize newly available habitat patches resulting from disturbance (Clobert et al. 2009, Reigada et al. 2015).

In early successional species, one might expect individuals to adjust their emigration and immigration decisions based on two main factors that potentially have an influence on reproductive success. First, large patches often support abundant resources and/or enhance the possibility of mate encounters, and therefore provide high fitness prospects. Emigration and immigration probabilities would thus be expected to be negatively correlated to patch size (Wahlberg et al. 2002, Schtickzelle & Baguette 2003, Gascoigne et al. 2009). Second, an early successional habitat patch only persists for a limited amount of time before it becomes unsuitable for breeding through the succession process (Turner et al. 1998, Prach & Walker 2011). The quality of the patch, and therefore the fitness prospects of early successional organisms, would be expected to decline over time (Duckworth 2012). Hence, when a patch is regularly disturbed (partly stopping the succession process), one might expect a negative relationship between the emigration and immigration probabilities and the extent of the patch’s surface area disturbed. Yet to our knowledge, few studies have investigated how patch-dependent fitness prospects may predict dispersal decisions in early successional organisms.

To study this issue, pond-breeding amphibians are suitable biological models, as many of these species reproduce in early successional aquatic habitats (Morand & Joly 1995, Cromer et al. 2002, Warren & Büttner 2008, Canessa et al. 2013). In temperate forests of Europe and North America, amphibians reproduce in temporary waterbodies, which in the past mainly resulted from natural disturbances (e.g. flooding or trees uprooted by wind, Joly & Morand 1994, DeMaynadier & Hunter 1995). Over the last century, however, human activity and forest management practices have rapidly modified the characteristics of amphibians’ aquatic breeding habitats, drastically affecting their population dynamics (see, for instance, Cayuela et al. 2016a, 2016b). The promotion of dense, monospecific forests with trees of uniform age has reduced the probability of windfall trees, decreasing the availability of naturally formed breeding waterbodies. In parallel, forest harvesting has led to the creation of semi-natural waterbodies (i.e. puddles formed in ruts and residual tracks made by skidders), which are now used by amphibians as replacement breeding habitats (Cromer et al. 2002, DiMauro & Hunter 2002, Kopecký et al. 2010). Yet these breeding habitats only persist for a limited time before they become unsuitable for reproduction due to natural silting. Accordingly, the long-term persistence of a local population strongly depends on (1) the continuous creation of new breeding patches (i.e. groups of ruts) that can be colonized to compensate for deterministic local extinctions triggered by waterbody silting dynamics and/or (2) frequent anthropic disturbance (i.e. from the passage of vehicles such as skidders) in existing patches to limit the natural silting process of waterbodies.

In this study, we examined how patch size and the level of patch disturbance affect reproductive success and breeding dispersal in a spatially structured amphibian population, the yellow-bellied toad (*Bombina variegata*). In forests exploited for timber, *B. variegata* breeds in early successional patches composed of waterbodies (e.g. ruts and residual puddles) resulting from logging operations (Cayuela et al. 2015). First, we investigated how patch size affects the probability of breeding occurrence and the abundance of newly metamorphosed individuals (i.e. local juvenile production) between patches. As large breeding patches usually increase the chance of mate encounter (Gascoigne et al. 2009) and provide more breeding resources (Cushman 2006), we predicted a positive relationship between breeding occurrence (i.e. presence of breeding indices), breeding success (i.e. presence and abundance of newly metamorphosed individuals) and patch size. In parallel, we also examined how the annual level of patch renewal through disturbance influenced breeding occurrence and success. By deepening ruts and increasing ground compaction (Wronski & Murphy 1994, Ampoorter et al. 2010), the passage of skidders limits the natural silting of waterbodies and improves their water-holding capacity, which reduces the risk of a pond drying out and thus amphibian reproductive failure (Tournier et al. 2017). Hence, we expected a positive relationship between breeding occurrence and success and the extent of the surface area disturbed in a patch each year by skidders. Secondly, we analyzed how patch size and disturbance influence breeding dispersal between patches. We took advantage of recent developments in multievent capture–recapture models (Cayuela et al. 2017a, Cayuela et al. 2018) to examine this issue. As individuals are expected to adjust their dispersal decisions according to their fitness prospects in a given patch, we hypothesized that emigration probability would be lowest in large patches with a higher disturbance level where reproductive success is highest. For the same reason, we hypothesized that immigration to these patches would also be highest.

## Materials and methods

### Study area and data collection

The capture–recapture (CR) study was conducted over a 9-year period (2000–2008) on a spatially structured population of *B. variegata* located in a woodland of northeast France (49.37°N, 4.83°E, elevation 200 m) (**Fig.1**). The woodland is a mixed forest covering approximately 7,000 ha and is surrounded by more or less intensively cultivated farmland. The nearest *B. variegata* population to our study population is separated by a distance of more than 20 km from the forest. The study population occupies 28 breeding patches (i.e. groups of ruts and puddles) in the study area. All the patches used by toads to reproduce were exhaustively sampled. The delineation of each patch was established using ArcGis 10.1 (ArcGis 10.1, Environmental Systems Research Institute, Redlands, California, USA) to create polygons connecting the waterbodies located on the boundary of the pond network (i.e. the Minimum Convex Polygon approach; see White & Garrott 1991). As in previous studies of *B. variegata* (see Cayuela et al. 2016a), patches were assumed to differ based on a minimum distance of 100 m separating the boundaries of polygons. This distance was chosen since the between-pond movement in this species is usually less than 100 m (Beshkov & Jameson 1980, Hartel 2008). The median of the Euclidean distance between patches was 2,458.83 m (min = 150.90 m, max = 6,905.91 m). Patch size was calculated as the mean cumulative surface area of the ruts composing the patch (i.e. length × width of each rut) over the 9-year study period. The surface area of the 28 breeding patches varied widely, ranging from 2.50 m^2^ to 107.67 m^2^ (mean = 23.99 m^2^, standard deviation = 22.33 m^2^). The patch disturbance level was evaluated by calculating the cumulative surface area of waterbody disturbed by skidder passages during a breeding season divided by the patch’s total water surface area (i.e. the percentage of the patch’s total water surface area that was disturbed). This variable was recorded only during the last two years of the study. For that reason, the capture–recapture analyses related to the effect of patch renewal through disturbance were restricted to the 2007–2008 period. The proportion of the waterbody surface area disturbed by skidders in the 28 breeding patches varied widely, from 0% to 100% (2007: mean = 47%, standard deviation = 46%, 2008: mean = 41%, standard deviation = 47%). Prior to the analyses, we verified the collinearity between patch size and disturbance; the two variables were weakly correlated (*r* = 0.15). We also examined the correlation between patch size and the number of waterbodies in a patch; these two variables were highly correlated (*r* = 0.73). Furthermore, we investigated how the number of adults in attendance in a breeding patch was related to patch characteristics (the analysis is detailed in **Appendix 1**). Zero-inflated Poisson regression models revealed that the number of adults increased with the size of the patch and the level of disturbance.

The capture sessions were carried out during the toad’s breeding season (from late April to July) of each year. The number of capture sessions per year ranged from 1 in 2003 and 2004 to 6 in 2000 (for details about the number of capture sessions and the number of individuals captured each year, see **Appendix 1**). During each capture session, all the breeding patches were exhaustively surveyed. Toads were caught by hand or using a dipnet. The catching effort was performed in a waterbody several times, and stopped when no new individual was detected. Based on the outcomes of previous studies on different *B. variegata* populations (Cayuela et al. 2016b), we assumed that toads became sexually mature at the age of 3, with a mean body length (snout–vent length) of 36 mm in males and 37 mm in females; smaller individuals were excluded from the analysis. The sex was assessed on the basis of strong forearms with nuptial pads in males. We identified each individual by the specific pattern of black and yellow mottles on its belly, recorded by photographs.

We surveyed breeding occurrence (i.e. presence of breeding indices) and success (i.e. presence and abundance of newly metamorphosed individuals) in 2007 and 2008. Two sampling sessions were performed annually in July and August to detect eggs, larvae and newly metamorphosed individuals.

This sampling period was chosen to coincide with the period during which larvae and newly metamorphosed individuals occur. The searches for eggs, larvae and newly metamorphosed individuals were conducted during the day by visual encounter. The occurrence of eggs and tadpoles and the number of newly metamorphosed individuals were recorded.

**Fig.1.**
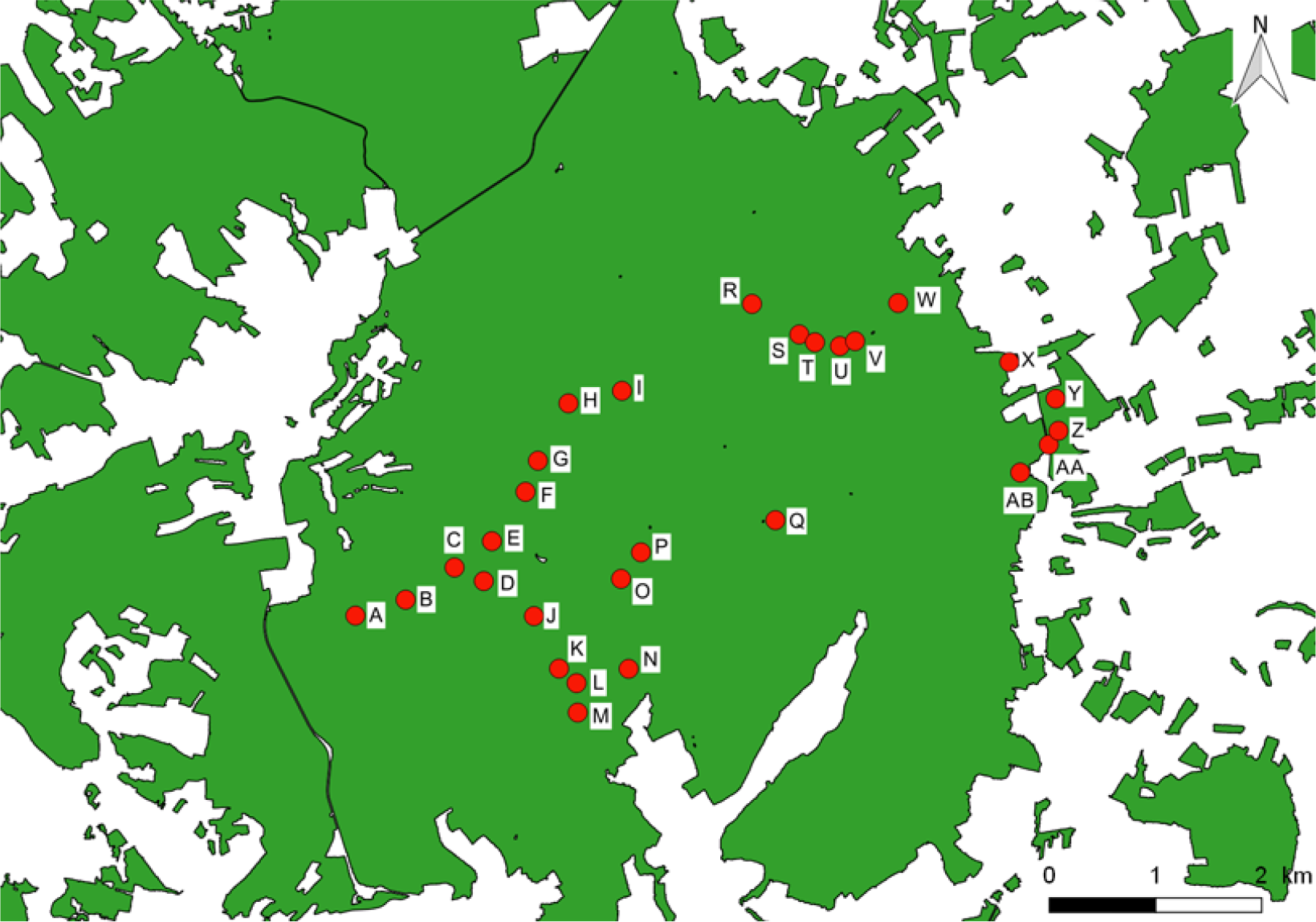
Map of the study area, showing the location (red dots) of the 28 breeding patches (i.e. group of ruts) within the woodland.

### Modeling the influence of patch size and disturbance on breeding occurrence and success

To investigate the influence of patch size and disturbance level on breeding occurrence, we used multistate occupancy models (Nichols et al. 2007, Gimenez et al. 2014). The data from the relevant two years of the breeding survey (2007 and 2008) was compiled in the same one dataset; one site therefore has two annual replicates with two detection occasions (July and August) each. The year was included in the model as a group effect. We considered three states: a site could be unoccupied for breeding (U), occupied with non-effective reproduction (i.e. presence of eggs and larvae only: L) or occupied with successful reproduction (i.e. presence of newly metamorphosed individuals: M). The observations were coded as undetected (0), eggs and/or larvae detected (1) and newly metamorphosed individuals detected (2). Following Gimenez et al. (2014), the model was based on three types of information (the model matrices are provided in **Appendix 1**). (1) The vector of initial state probabilities contained two parameters of interest: the probability that a site is occupied with non-effective reproduction (ψ1), and the probability that a site is occupied with successful reproduction (ψ2). (2) The state–state transition matrix contained the transition probabilities, which were fixed at 1 in our study case; we did not model changes in the state of occupancy over time. (3) The field observation matrices allowed us to model the observation process conditional on underlying occupancy states. Two modeling steps were considered to highlight the successive processes of detection and breeding-state ascertainment. In the first matrix (**Appendix 1**), we introduced a set of intermediate observations: undetected (*u*), detection of eggs and larvae (*l*), and detection of newly metamorphosed individuals (*m*). This resulted in the consideration of two detection probabilities: the detection probability of eggs and/or larvae (*p*1), and the detection probability of newly metamorphosed individuals (*p*2). The second matrix (**Appendix 1**) specified the probability of successful reproduction conditional on these intermediate observations. This parameterization was implemented in the E-Surge program (Choquet et al. 2009). We ranked the models using Akaike information criteria adjusted for a small sample size (AICc) and AICc weights. When the AICc weight of the best-supported model was less than 0.9, we performed model averaging. We tested our hypotheses from the general model, [ψ1,2(Di + Si), *p*1,2(Y)], which included three variables: the proportion of the surface area of a waterbody disturbed by skidders (Di, a continuous variable), the size of the patch (Si, a continuous variable), and a year effect (Y, a discrete variable with two modalities, 2007 and 2008). Due to reduced statistical power (30 patches sampled over a 2-year period), the two patch-specific variables were included in an additive way in the model. For the same reason, we also considered an additive effect between two conditional occurrence probabilities (ψ1 and ψ2) and the variables. We tested all combinations of these variables, leading to the consideration of 8 competing models (**Appendix 1**).

Then we conducted a second analysis to examine how patch size and disturbance influenced the number of individuals that successfully metamorphosed in the breeding patch. As newly metamorphosed individuals cannot be marked using a non-invasive method and leave the pond shortly after metamorphosis, we could not use density estimates based on capture–recapture or repeated count data. As the detection probability of newly metamorphosed individuals estimated by our multistate occupancy model was high (> 0.90, see ‘Results’), we assumed that imperfect detection would not skew our inferences. To avoid the risk of double counting, we considered in our analyses the maximum number of newly metamorphosed individuals recorded in each patch during one of the two annual sampling sessions. Due to the zero excess in the count data, we used zero-inflated Poisson (ZIP) regression models. First, we performed a preliminary analysis to investigate how the number of newly metamorphosed individuals was correlated with the number of adults in each patch and the density of adults per m^2^ of patch. The number and the density of adults were corrected by dividing the number of captured individuals by the recapture probability (i.e. Horvitz-Thompson estimator) estimated by CR multievent models (see below): 0.45 in 2007 and 0.46 in 2008. The number of newly metamorphosed individuals (Ju) was treated as the response variable. In the Poisson regression part of the model, the number of adults (Ad), the density of adults (De), and the year (Y: 2007, 2008) were introduced as explanatory terms, resulting in the following general model (Ju ~ Ad + De + Y). The variables (Ad) and (De) were treated as continuous variables, were z-scored and were included in an additive way. We examined all the possible combinations of effects, resulting in 8 competing models. The models were ranked using Akaike information criteria adjusted for a small sample size (AlCc) and AlCc weights. Normality of the residuals of the best-supported model was examined graphically using a quantile–quantile plot. Then we examined how newly metamorphosed individuals across patches (Ju) was correlated to patch size (Si) and disturbance (Di). We used the same modeling approach (i.e. ZIP models) as described above. The variables (Si) and (Di) and the year (Y: 2007, 2008) were incorporated as explanatory terms. The continuous variables (Si) and (Di) were z-scored. Moreover, as our preliminary analysis showed that the response variable (Ju) was strongly correlated with the number of adults (Ad) in patches, we introduced the variable (Ad) as an offset, leading to the following general model: [Ju ~ offset(log(1+Ad)) + Di + Si + Y]. From this model, we examined all possible combinations of effects.

**Table 1.**
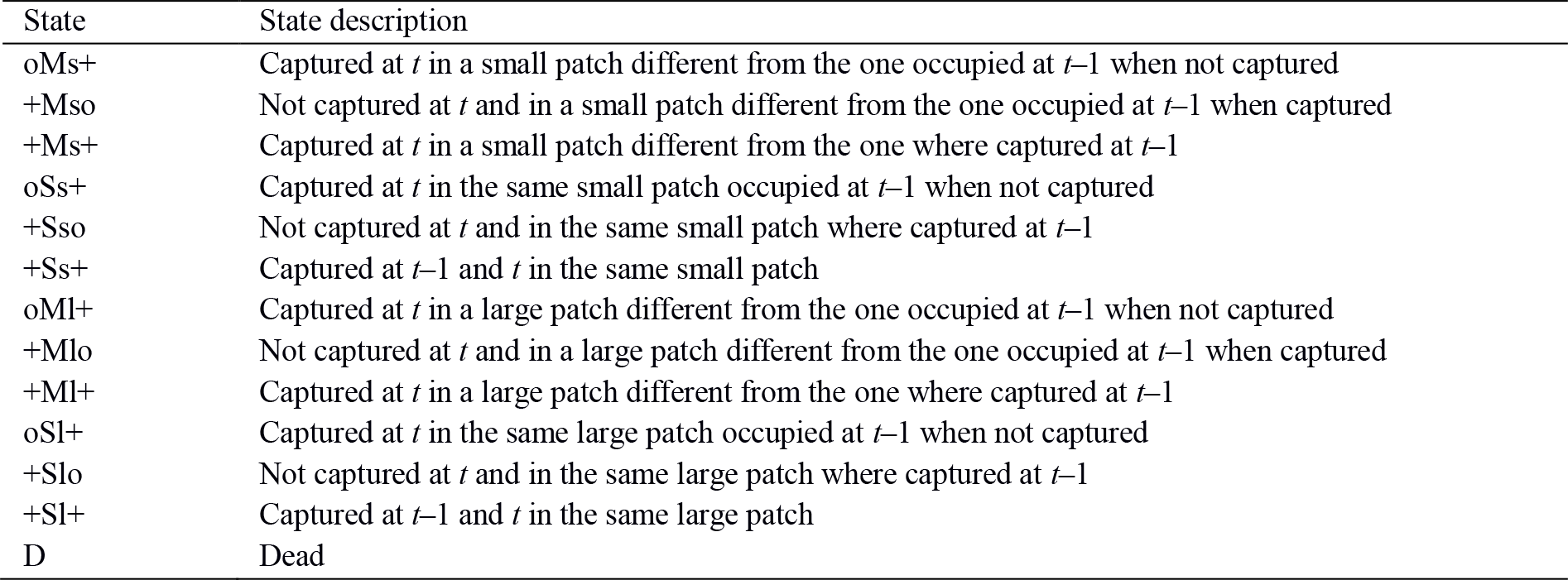
The 13 states of the CR multievent models and their definitions. The state formulation includes four types of information: + = captured or o = not captured at previous occasion, S = stayed or M = moved, s = small patch or l = large patch, + = captured or o = not captured at current occasion. Note that the patch disturbance model used the same states as those presented here except that ‘s’ (small) and ‘l’ (large) were replaced by l = low disturbance and h = high disturbance.

**Fig.2.**
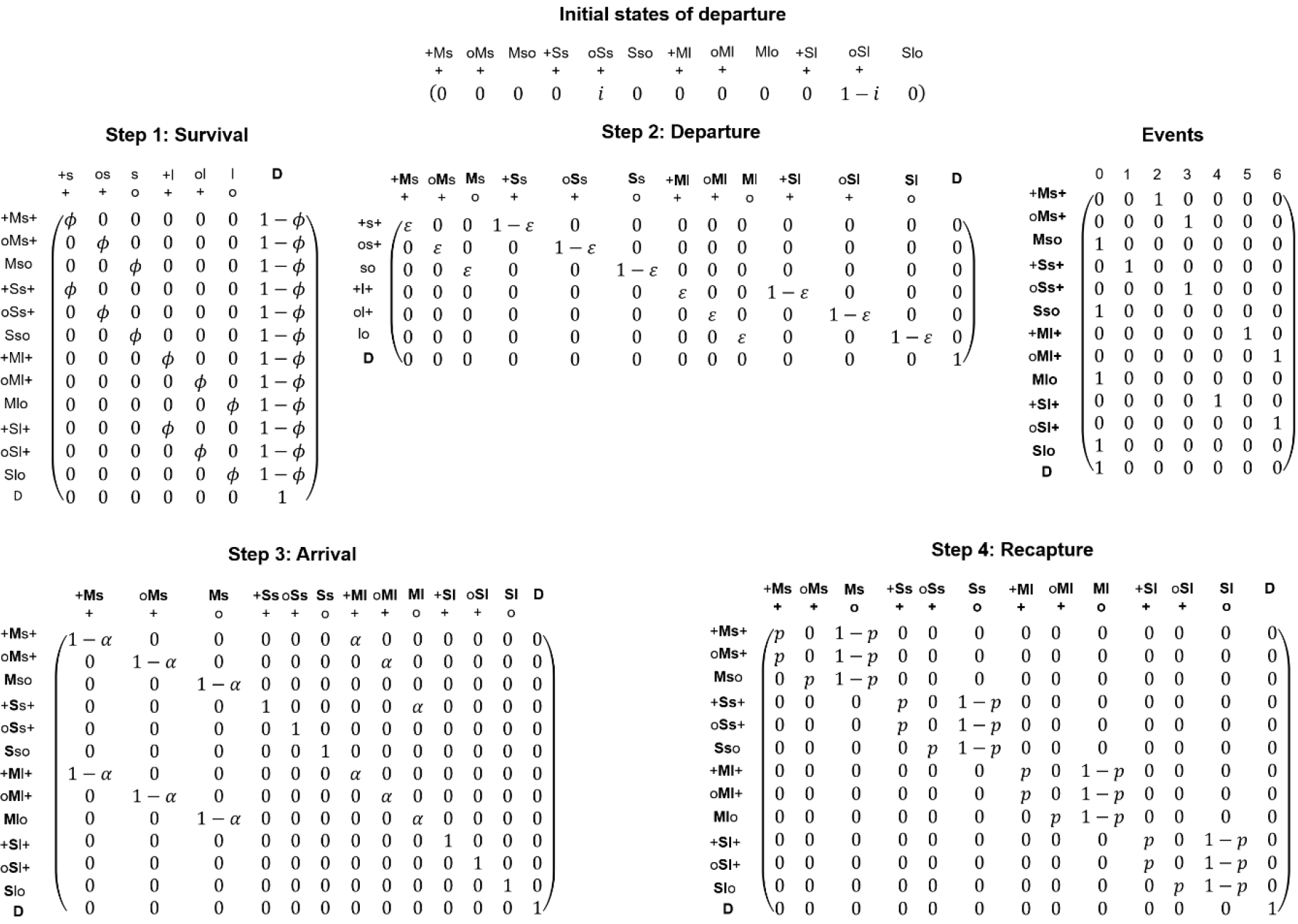
Modeling the influence of patch size: matrices of initial states of departure, state–state transitions and events (field observations). The definition of the states is provided in Table 1. Four state–state transition steps were considered in the model: survival probability ϕ (step 1), departure probability ψ (step 2), arrival probability α (step 3) and recapture probability p (step 4).

### Modeling the influence of patch size on dispersal

To test the effect of patch size on dispersal, we used CR multievent models recently developed by Cayuela et al. (2017a) and further extended by Tournier et al. (2017), which allow the estimation of movement rates between sites with differing characteristics. As with other multievent models, this model is composed of states and events (Pradel 2005). Events correspond to field observations and are coded in an individual’s capture history. These observations are related to the latent state of the individuals. Yet observations can carry a certain degree of uncertainty regarding this latent state. Multievent models aim to model this uncertainty in the observation process using hidden Markov chains.

The model used in Tournier et al. (2017) is based on 13 states (**Table 1**) that combine information about whether or not an individual occupies the same patch as on the previous capture occasion, information about whether or not the individual was/is captured at the previous and current occasion, and on the category of patch currently occupied. Note that in CR multievent models, patch characteristics cannot be introduced as continuous variables and are treated in a discrete way (Cayuela et al. 2017a, 2018). The following codes are used: an individual that occupies the same patch as on the previous occasion is coded S for ‘stayed’ or if it occupies a different patch, M for ‘moved’. These codes are prefixed by the previous capture status and suffixed by the current capture status (+ for ‘captured’, o for ‘not captured’). In addition, we added a state designation to include the category of patch currently occupied: an individual can occupy one of two patch categories (*s* ‘small’ or *l* ‘large’). A small patch ranged from 2.5 m^2^ to 23 m^2^, and a large patch ranged from 23 m^2^ to 108 m^2^. We used the mean size of patches in our study system as the boundary between the two size classes. Five events were considered in the model: for an individual captured at *t* and *t*–1, a code of 1 or 4 was attributed if it did not change patch and was in patch category *s* or *l* respectively, and a code of 2 or 5 was attributed if it did change patch and was in patch category *s* or *l* respectively. For an individual not captured at *t*–1 and captured at *t* in patch category *s* or *l*, a code of 3 or 6 was attributed respectively. An individual not captured at *t* was given a code of 0. The model had a robust design structure (Pollock 1982), i.e. several capture sessions performed within a year corresponded to secondary sessions, and a set of yearly sessions corresponded to a primary period. This robust design structure allowed us to examine both intra-annual and inter-annual dispersal.

At its first capture, an individual could be in state oSs+ or oSl+. From this initial state of departure, the transition from the state at time *t*–1 to that at time *t* was updated through four successive modeling steps: (1) survival, (2) departure, (3) arrival, and finally, (4) recaptured or not (see **Fig.2**). Following the convention set out in Souchay et al. (2014), whenever the status in the state descriptor was updated to the situation at *t*, it became bold (and stayed bold throughout the following steps). First, the information about survival was updated; an individual could survive with a probability of ϕ, or die with a probability of 1–ϕ (**Fig.2**), leading to a transition matrix with 13 possible states of departure and 7 intermediate arrival states. Survival probability was set at one between secondary sessions. Second, departure was updated; an individual that survived between *t*–1 and *t* could leave its patch (designated *s* or *l*) with a probability of ψ or stay in the same patch with a probability of 1–ψ (**Fig.2**). The departure probability could be dependent on the category of the patch (*s* or *l*). This resulted in a transition matrix with 7 intermediate departure and 13 intermediate arrival states. Third, arrival was updated; an individual that left its patch could arrive in a small patch (*s*) with a probability of 1–α or in a large patch (*l*) with a probability of α (**Fig.2**). This led to a transition matrix with 13 intermediate departure states and 13 intermediate arrival states. Fourth, the recapture status of individuals was updated; an individual could either be captured with a probability of *p* or not captured with a probability of 1–*p* (**Fig.2**), resulting in a transition matrix with 13 intermediate departure states and 13 arrival states at time *t*. The last component of the model linked events to states. In this specific situation, each state corresponded to only one possible event (**Fig. 2**).

The parameterization was implemented in the E-Surge program (Choquet et al. 2009). As the information about patch size was recorded over the entire 9 years of the survey, the models were run using the complete dataset (2000-2008). Competing models were ranked using Akaike information criteria adjusted for a small sample size (AlCc) and AlCc weights. When the AICc weight of the best-supported model was less than 0.9, we performed model averaging. The 95% CIs of model-averaged parameters were calculated using the parametric bootstrap method. Our hypotheses regarding recapture and state–state transition probabilities were examined using the general model [ϕ(Si + S), ψ(Si + S), α(Si + S), *p*(Y + Si + S)], which included three effects: (1) patch size (Si), coded as states in the model; (2) group effect for sex-specific variation (S); and (3) year-specific variation (Y). As previous studies on this toad population have shown that recapture probability varies between years (Cayuela et al. 2016a, 2016b), we retained year-specific variation in all the models. From this general model, we tested all the possible combinations of effects, resulting in the consideration of 64 competing models (**Appendix 2**).

### Modeling the influence of patch disturbance on dispersal

The effect of patch disturbance was evaluated using the model recently proposed by Cayuela et al. (2018) to account for the possibility that a patch could change category (i.e. low ↔ high disturbance) between two capture occasions. We used the mean proportion of disturbed surface area as the boundary between the two classes. Low disturbance corresponded to patches with a lower proportion of the waterbody surface area disturbed by skidder passages (from 0% to 41%, the latter being the mean disturbance rate). High disturbance corresponded to patches with a higher proportion of the waterbody surface area disturbed by skidders (from 42% to 100%). This model was based on the same states (**Table 1**) and events as the previously described model; 13 states and 7 events were thus considered. In contrast to the previous model, we also considered additional state–state transitions in the arrival matrix to update patch status (step 3, **Fig.3**). In addition, we included the code *l* for ‘low disturbance to patch’ and *h* for ‘high disturbance to patch’. The model had a robust design structure to examine both intra-annual and interannual dispersal.

At its first capture, an individual could be in state oSl+ or oSh+. Then the transition from the state at time *t*–1 to that at time *t* was updated through four successive modeling steps: (1) departure, (2) survival, (3) arrival and patch dynamics, and, finally, (4) recaptured or not (**Fig.3**). First, information about departure was modeled; an individual could move from a patch (designated *l* or *h*) with a probability of ψ or remain in the same patch occupied at *t*–1 with a probability of 1–ψ. The departure could be made dependent on the category of patch occupied at *t*–1. This led to a transition matrix with 13 states of departure and 9 intermediate states of arrival (**Fig.3**). Second, survival was updated; an individual could survive with a probability of ϕ or die with a probability of 1–ϕ, resulting in a transition matrix with 9 possible states of departure and 9 arrival states. Third, arrival and site dynamics were modeled. When an individual remained in the same site, it could occupy a site that changed state (*l* ↔ *h*) between *t*–1 and *t* with a probability of α, or a site that remained in the same state with a probability of 1–α (**Fig.3**). When an individual moved, it could arrive in a site of a different category (*l* or *h*) than the one previously occupied with a probability of α or in the same type of site with a probability of 1-α (**Fig. 3**). This led to the consideration of a transition matrix with 9 departure states and 9 arrival states (**Fig.3**). Fourth, recapture status was updated; an individual could be recaptured with a probability of *p* or missed with a probability of 1-*p*. Recapture probability could depend on the patch category, leading to the consideration of a transition matrix including 9 departure states and 13 arrival states (**Fig.3**). The last component of the model linked events to states (**Fig.3**).

**Fig.3.**
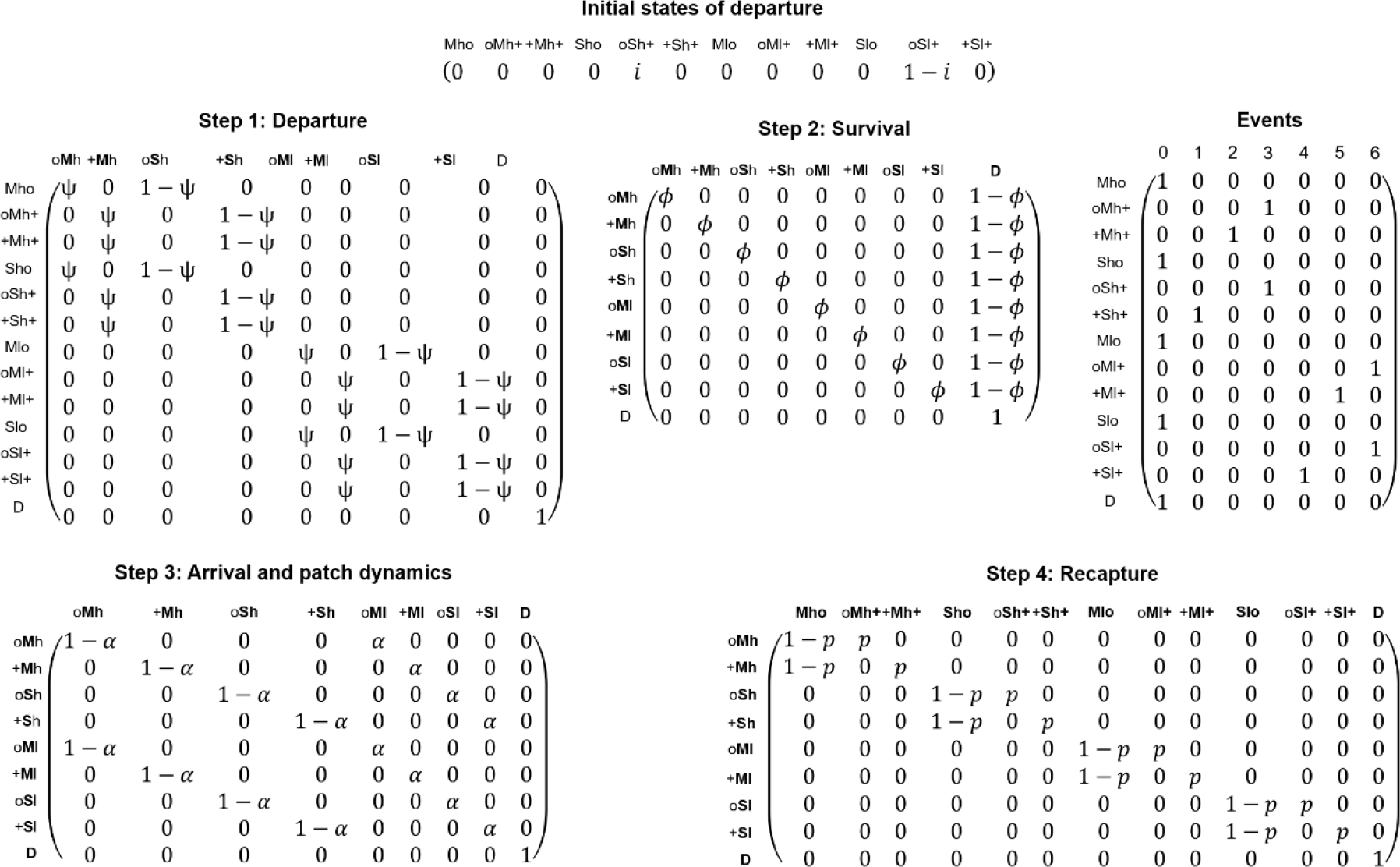
Modeling the influence of patch disturbance: matrices of initial states of departure, state–state transitions and events (field observations). The definition of the states is provided in Table 1. Four state–state transition steps were considered in the model: departure probability ψ (step 1), survival probability ϕ (step 2), probability of arrival and patch state change α (step 3) and recapture probability *p* (step 4).

The parameterization was implemented in the E-Surge program. As the information about patch disturbance was only recorded during the last two years of the survey, in this case the models were run using only the 2007–2008 dataset. Competing models were ranked using Akaike information criteria adjusted for a small sample size (AICc) and AICc weights. We performed model averaging when the AICc weight of the best-supported model was less than 0.9 and used the parametric bootstrap method to calculate the 95% CI. Our hypotheses about departure ψ, survival ϕ, arrival α and recapture *p* were examined using the general model [ψ(Di + S), ϕ(S), α(Di + S), *p*(Y + Di + S)], which included three effects: (1) the patch disturbance level (Di), coded as states in the model; (2) group effect for sex-specific variation (S); and (3) year-specific variation (Y). As this analysis was based on a subset of data including only two years of study, we did not test the effect of disturbance on survival due to a lack of power. In all the models, the probability that a patch changed disturbance level depended on the site status at *t*–1. From the general model, we tested all the possible combinations of effects, resulting in the consideration of 32 competing models (**Appendix 2**).

### Assessing the effects of patch size and disturbance on arrival probability

The conditional probability of arrival (i.e. depending on patch characteristics) estimated by the multievent CR models strongly depended on the quantity of patches of each category in the spatially structured population and the number of individuals in each patch that may reach these. For this reason, we examined patch size and disturbance by comparing the model-averaged conditional arrival probability to the probability of reaching a patch using a random dispersal hypothesis (i.e. the mean probability of arriving in a patch calculated from all the individuals occurring in all patches of the study area). As the number of males and females varied between patches, we calculated the expected random sex-specific probability. We assumed that the effect of patch size or disturbance was significant if the 95% CI of the conditional arrival probability did not overlap with the expected random probability. The percentage of deviation from the expected random probability was used to assess the influence of patch characteristics and to rank their effects on both sexes.

## Results

### Influence of patch size and disturbance on breeding occurrence and reproductive success

Multistate models revealed that the occurrence probability of non-effective reproduction was 0.10 (95% CI 0.04–0.23), whereas the probability of successful reproduction was 0.40 (95% CI 0.27–0.55); these estimates were extracted from a model with constant occurrence probability [ψ1,2(.), *p*1,2(Y)]. The detection probability of breeding indices (eggs and tadpoles) was 0.96 (95% CI 0.63–0.99); for newly metamorphosed individuals it was 0.93 (95% CI 0.83–0.97). The best-supported model was [ψ1,2(Si), *p*(Y)]; the complete model selection procedure is provided in **Appendix 1**. As its AICc weight was 0.58, we performed model averaging. Detection probability of non-effective reproduction was 0.96 (95% CI 0.60–0.99) in 2007 and 0.97 (95% CI 0.54–0.99) in 2008. Detection probability of successful reproduction was 0.91 (95% CI 0.76–0.97) in 2007 and 0.94 (95% CI 0.79–0.98) in 2008. More importantly, our analysis revealed that both non-effective and successful breeding probabilities were positively influenced by patch size (model-averaged slope: 3.00, 95% CI 0.71–5.41), whereas disturbance level had a marginal effect (model-averaged slope: 0.08, 95% CI −0.15–1.03).

The number of newly metamorphosed individuals in 2007 and 2008 varied from 0 to 73 from one patch to another (mean = 5.33, sd = 14.60). Our preliminary analysis revealed that the number and density of adults influenced the number of metamorphosed individuals in a patch. The best-supported ZIP model (Ju ~ Ad + De + Y) had an AICc weight of 0.99 (**Appendix 1**). It showed that the number of newly metamorphosed individuals was positively influenced by the number of adults recorded in the patch and negatively affected by the density of adults per m^2^ of the patch (**Fig. 4A** and **4B**). We then analyzed the effects of patch size and disturbance on the number of newly metamorphosed individuals, taking into account the number of adults in the patch. The best-supported ZIP model [Ju ~ offset(log(1+Ad)) + Di + Si + Y] had an AICc weight of 0.99 (see the complete model procedure in **Appendix 1**). The number of newly metamorphosed individuals was positively influenced by both patch size and patch disturbance (**Fig. 4C** and **4D**). The number of newly metamorphosed individuals was also higher in 2008 than in 2007 (**Fig. 4C** and **4D**).

**Fig.4.**
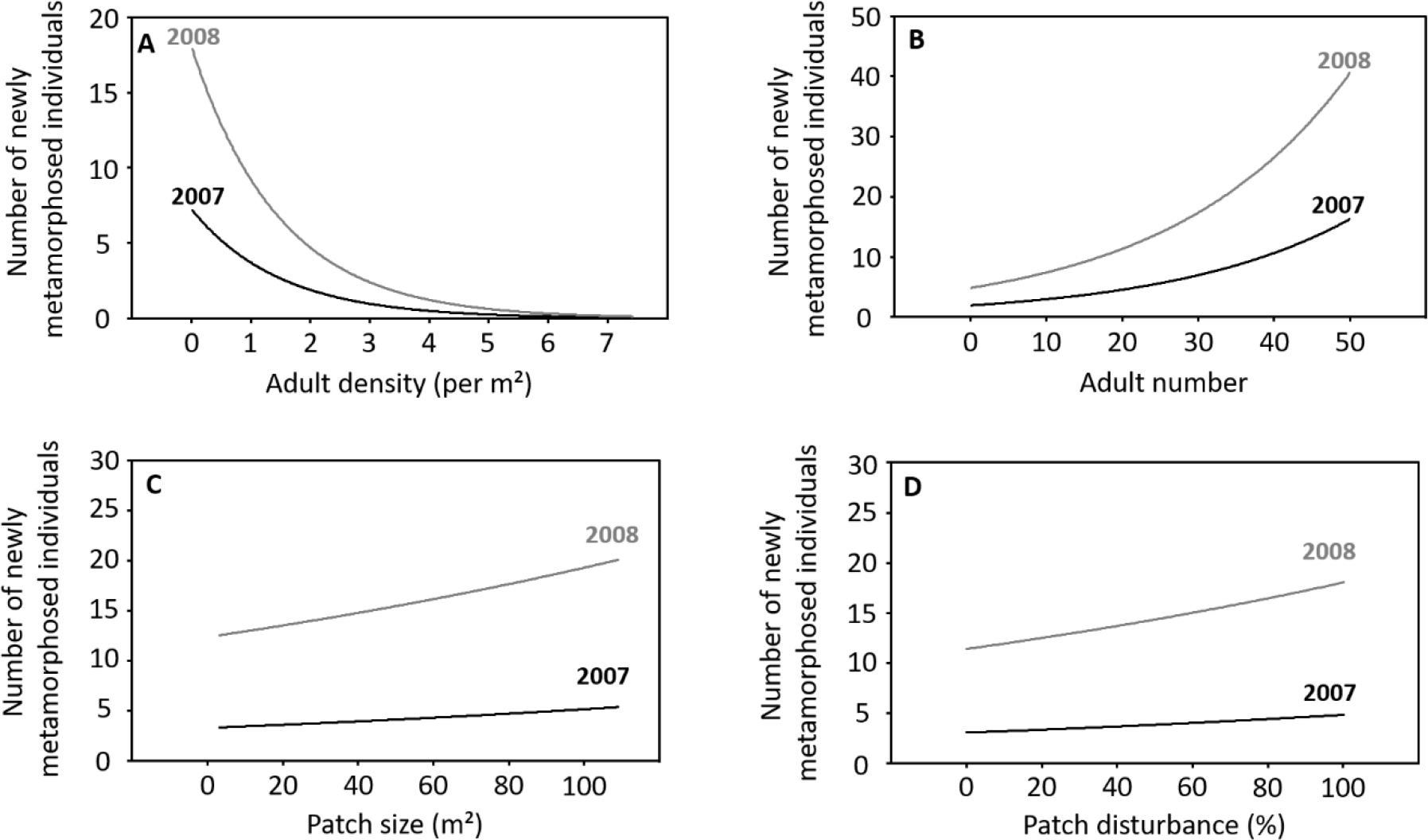
Influence of the number and the density of adults, patch size, and the level of disturbance on the breeding success of the yellow-bellied toad (Bombina variegata), expressed by the number of newly metamorphosed individuals (in 2007 and 2008). The outputs of zero-inflated Poisson regression models are shown. While the number of adults in attendance in a patch increased the number of newly metamorphosed individuals (B), the density of adults exerted an opposite effect (A). The number of newly metamorphosed individuals increased with patch size (C) and the proportion of the patch’s surface area disturbed by skidders (D).

### Influence of patch size and disturbance on dispersal

During the 9-year study period, we captured 744 toads. Of these, we identified 230 adults (120 males and 110 females). The number of individuals captured each year varied from 14 in 2004 to 103 in 2007 (for more details, see **Appendix 2**). We detected 64 dispersal events between successive capture sessions.

Concerning the influence of patch size on dispersal, the best-supported model was [ϕ(.), ψ(Si), α(Si), *p*(Si)]; the complete model selection procedure is provided in **Appendix 2**. As the AICc weight of this model was 0.13, we performed model averaging. Recapture varied according to patch size and between years: recapture probability was higher in small patches than in large ones. In small patches, recapture probability varied from 0.23 (95% CI 0.16–0.32) in 2003 to 0.59 (95% CI 0.51–0.66) in 2000, while in large patches it varied from 0.18 (95% CI 0.13–0.24) in 2003 to 0.50 (95% CI 0.44–0.57) in 2000. Sex-specific variation in recapture was marginal. In terms of survival, model-averaged estimates indicated that this did not vary according to patch size. Rather, the estimates suggested weak sex-specific variation in survival: males had a slightly lower survival probability (0.78, 95% CI 0.70–0.81) than females (0.80, 95% CI 0.76–0.86). More importantly, the analyses showed that dispersal depended on patch size (**Fig.5A**). At both intra- and inter-annual levels, individuals had a higher probability of leaving a small patch than a large one. Intra-annually, the probability of an individual leaving a small patch was 0.19 (95% CI 0.14–0.27) in males and 0.16 (95% CI 0.10–0.21) in females; in contrast, in large patches it was 0.09 (95% CI 0.07–0.15) in males and 0.08 (95% CI 0.05–0.11) in females. We found the same pattern at the inter-annual level (**Fig.5**). Furthermore, arrival probability also depended on patch size (**Fig.5B**). Intra-annually, the proportion of individuals arriving in a large patch was 0.62 (95% CI 0.48–0.75) in males and 0.58 (95% CI 0.40–0.71) in females. At the inter-annual level, we detected a similar pattern: the proportion of individuals arriving in a large patch was 0.49 (95% CI 0.34–0.68) in males and 0.46 (95% CI 0.28–0.61) in females. These values were systematically far higher than those expected based on a random dispersal hypothesis (0.32 in males and females). The deviation of the estimated from the expected value was 94% in males and 81% in females at the intra-annual level; inter-annually, the deviation was 53% in males and 44% in females.

Concerning the influence of patch disturbance, the best-supported model was [ψ(Di), ϕ(S), α(Di), p(Y)]; the complete model-selection procedure is provided in **Appendix 2**. As the AICc weight of this model was 0.24, we performed model averaging. The model-averaged estimates indicate that dispersal depended on the level of patch disturbance (**Fig.5C**). Intra-annually, the probability of an individual leaving a patch with high disturbance was 0.03 (95% CI 0.01–0.10) in both males and females; in contrast, in patches with low disturbance it was 0.26 (95% CI 0.15–0.40) for both sexes. A similar pattern was detected at the inter-annual level. In contrast, arrival probability did not vary significantly according to patch disturbance (**Fig.5D**). The 95% CI overlapped with the expected probability under random dispersal hypothesis. However, it is probable that this result is due to a lack of statistical power since the 95% CI was large (likely because we used a small subset for our analyses), while the deviation between estimated probability and expected probability (0.44 in males and females) was wide. Intra-annually, the proportion of individuals arriving in a patch with high disturbance was 0.71 (95% CI 0.44–0.88) in males and females. At the inter-annual level, the proportion of individuals arriving in a patch with high disturbance was 0.79 (95% CI 0.45–0.95) in both sexes. The deviation of the estimated from the expected value was 61% intra-annually; at the inter-annual level, the deviation was 80%.

**Fig.5.**
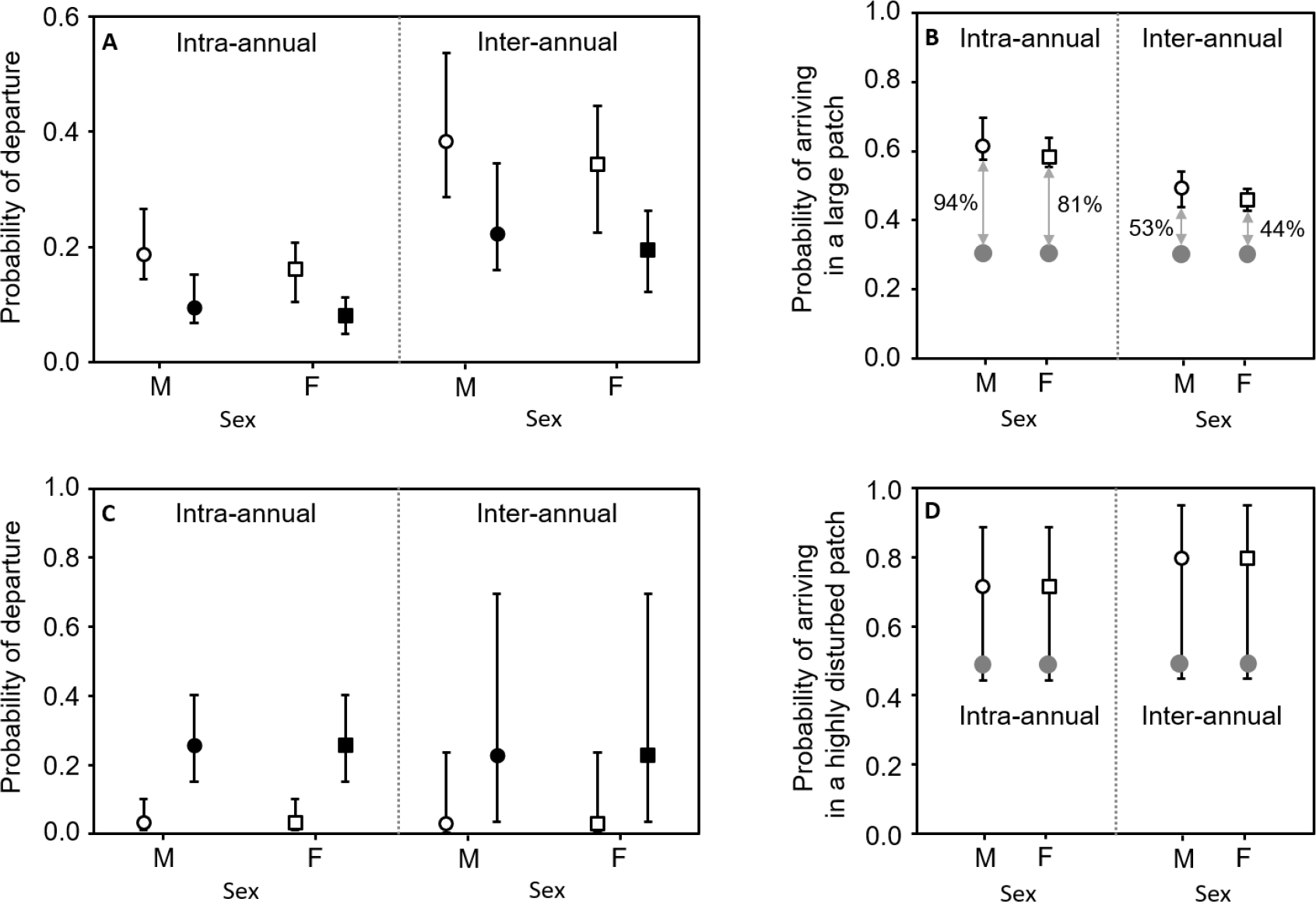
Influence of patch size and level of disturbance on dispersal in the yellow-bellied toad (Bombina variegata) intra-annually and inter-annually. (A) The departure probability of males (circles) and females (squares) depending on patch size (empty circles and squares = small patch; full circles and squares = large patch). Error bars show the 95% CI. (B) The probability of arriving in a large patch differed from the expected probability under a random dispersal hypothesis (grey circles); the 95% CI did not overlap the expected probability. The percentages correspond to the deviation between the estimated and expected probabilities. (C) The departure probability of males (circles) and females (squares) depending on the level of patch disturbance (empty circles and squares = patch with high disturbance; full circles and squares = patch with low disturbance). (D) The probability of arriving in a patch with high disturbance did not significantly differ from the expected probability under a random dispersal hypothesis (grey circles); the 95% CI overlapped the expected probability.

## Discussion

Our findings revealed that both the probability of successful reproduction of *B. variegata* and the number of newly metamorphosed individuals increased with patch size as well as with the level of disturbance. In addition, the results showed that the patch-specific factors affecting breeding success also influenced breeding dispersal. Emigration probability was negatively influenced by larger patch size; in contrast, larger patch size positively influenced immigration probability. Equally, a higher level of breeding patch disturbance (measured by the percentage of surface area of the patch disturbed each year) had a strong negative influence on emigration probability; simultaneously, a higher level of disturbance slightly positively affected immigration probability.

### Influence of patch size and disturbance on local breeding success and fitness prospects

The findings showed that the occurrence probability of newly metamorphosed individuals was higher in large patches than in small patches. We also found that the number of metamorphosed individuals increased with the number of adults in attendance in a breeding patch, which was positively correlated with patch size. Yet the analyses showed that the number of newly metamorphosed individuals still increased with patch size even after controlling for the number of adults in a patch. This indicates that patch size affects local breeding success regardless of breeder abundance. This might be explained in two ways: first, reproducing in a large patch with extensive breeding resources may reduce intraspecific larval competition and enhance breeding success. However, an increase in reproductive success remains conditional on the local population density, since our results found that the number of newly metamorphosed individuals decreased with the density of breeding adults per m^2^ (**Appendix 1**). A high density of adults in a patch likely enhances larval density and competition, which negatively affects tadpole growth and survival (Jasieński 1988). Second, large patches contain a higher number of breeding ponds than small ones. In *B. variegata,* females usually spread their egg capital over several clutches deposited in different waterbodies that often vary in temperature, hydroperiod and trophic resources (Barandun et al. 1997). This egg-spreading behavior is often regarded as a bet-hedging tactic to reduce the risk of reproduction failure in the face of hydric instability of waterbodies (Barandun et al. 1997, Buschmann 2002).

The probability of the occurrence and the number of newly metamorphosed individuals increased with the proportion of the surface area of the waterbody disturbed by skidders; this increase was nevertheless marginal to the probability of successful reproduction, which may be due to the intrinsic roughness of the occurrence data and the small number of breeding patches (28) in the study area. This result is congruent with previous studies highlighting that the occurrence of *B. variegata* reproduction is associated with aquatic habitats disturbed by human activity (Warren & Büttner 2008, Canessa et al. 2013). In particular, Warren & Büttner (2008) found that *B. variegata* preferentially occupied and bred in waterbodies whose ground surface area disturbed by vehicle passage ranged from 40-100%. By deepening ruts and enhancing ground compaction (Wronski & Murphy 1994, Ampoorter et al. 2010), skidders limit the natural silting of waterbodies and increase their water-holding capacities, thus decreasing their risk of drying out. By reproducing in patches that include waterbodies with a longer hydroperiod, individuals can therefore mitigate the risks of breeding failure caused by desiccation, which is the main mortality factor at larval stage (Barandun & Rever 1997, Barandun et al. 1997).

### Influence of patch size and disturbance on dispersal

We found that patch-specific factors influencing local breeding success also affected emigration and immigration probabilities. First, individuals were less likely to emigrate from large patches (where breeding success was highest), than from small ones; similarly, individuals were more likely to immigrate to large patches. Second, individuals had a lower probability of emigrating from patches experiencing a higher level of disturbance (defined by 4–100% of the patch’s surface area disturbed annually by skidders), where breeding success was highest, than from patches with a low level of disturbance (0–41% of the surface area disturbed). Our results also suggested that individuals were more likely to immigrate to patches with higher disturbance. These asymmetric dispersal rates indicate that individuals adjust their dispersal decisions according to breeding patch characteristics, likely basing their choices on environmental and/or social signals that provide valuable information about local fitness prospects. This conclusion is in accordance with a recent study showing that *B. variegata* individuals were less likely to leave waterbodies where the risk of drying out is low, thus favoring successful and constant reproduction over time (Tournier et al. 2017). In anurans, individuals use olfactory cues to locate ponds and evaluate their quality for breeding (Sinsch 1990, Semlitsch 2008), behavior that has also been observed in *B. variegata* (Cayuela et al. 2016c, 2017b). This behavioral mechanism likely allows individuals to assess their chances of breeding success in a patch and then to decide where to breed.

## Conclusion

The results of this study highlight that patch size and the level of disturbance affect the chances of reproductive success in *B. variegata.* They also suggest that breeders adjust their dispersal decisions according to local fitness prospects. In early successional organisms, a plastic response to dispersal likely permits rapid adjustment to progressive changes in environmental conditions resulting from the ecological succession process. This results in non-random dispersal between patches, which would be expected to have dramatic consequences on the demography of spatially structured populations by affecting local recruitment and population size. In addition, non-random dispersal is likely to drive the direction and intensity of gene flow, and could thus influence evolutionary processes in a patch by affecting the effective population size, the effects of genetic drift, and the effectiveness of selection. Future studies could help gain a better understanding of the eco-evolutionary dynamics shaping the demography and the evolution of early successional species.

## Acknowledgments

We warmly thank all the fieldworkers who assisted in data collection. This research project was funded by the Lorraine Direction Régionale de l’Environnement, de l’Aménagement et du Logement (DREAL), the Agence de l’Eau Rhin-Meuse, the Conseil Régional de Lorraine, the Conseil Régional de Champagne-Ardenne, the Conseil Régional de Picardie, the Conseil Général de l’Aisne, the Conseil Général d’Ardéche, the Conseil Général d’lsére and the Communauté de Communes de l’Argonne Ardennaise (2C2A).

Author contributions
JP, RH and PJ designed the capture–recapture survey. JP performed the survey and collected the data in the field. LB, AB and HC analyzed the data. LB and HC wrote the manuscript; all the authors provided editorial advice.

